# HexSE: Simulating evolution in overlapping reading frames

**DOI:** 10.1101/2022.09.09.453067

**Authors:** Laura Muñoz-Baena, Kaitlyn E. Wade, Art F. Y. Poon

**Affiliations:** Department of Microbiology and Immunology, Western University, London, ON, Canada; Department of Pathology and Laboratory Medicine, Western University, London, ON, Canada

## Abstract

**Motivation:** Gene overlap occurs when two or more genes are encoded by the same nucleotides. This phenomenon is found in all taxonomic domains, but is particularly common in viruses, where it may provide a mechanism to increase the information content of compact genomes. The presence of overlapping reading frames (OvRFs) can skew estimates of selection based on the rates of non-synonymous and synonymous substitutions, since a substitution that is synonymous in one reading frame may be non-synonymous in another, and vice versa.

**Results:** To understand the impact of OvRFs on molecular evolution, we implemented a versatile simulation model of nucleotide sequence evolution along a phylogeny with any distribution of open reading frames in linear or circular genomes. We use a custom data structure to track the substitution rates at every nucleotide site, which is determined by the stationary nucleotide frequencies, transition bias, and the distribution of selection biases (dN/dS) in the respective reading frames.

**Availability and implementation:** Our simulation model is implemented in the Python scripting language. All source code is released under the GNU General Public License (GPL) version 3, and is available at https://github.com/PoonLab/HexSE.

## INTRODUCTION

Overlapping reading frames (OvRFs) are portions of the genome where the same nucleotide sequence encodes more than one protein. They have been documented across all taxonomic domains including bacteria, vertebrates and fungi (Pallejà et al., 2008; Ribrioux et al., 2008; Chung et al., 2007; Gerads and Ernst, 1998). OvRFs are particularly abundant in virus genomes and examples can be found in all Baltimore classes (Muñoz-Baena and Poon, 2022). In viruses, OvRFs may provide a mechanism to store more information in smaller genomes, to make purifying selection more efficient at removing deleterious mutations, or to facilitate the *de novo* creation of genes (Willis and Masel, 2018; Sabath et al., 2012; Belshaw et al., 2008; Krakauer and Plotkin, 2002; Chirico et al., 2010). OvRFs can be classified into six different frameshifts that can be annotated as +2, + 1, +0, −0, −1, −2 (Lèbre and Gascuel, 2017), although other labeling schemes exist. Here, the sign indicates whether the overlap occurs between open reading frames (ORFs) on the same (+) or the opposite (−) strands, and the integer indicates how many nucleotides they are shifted relative to one another. Frameshifts influence the effect of selection in OvRFs. For example, a +0 frameshift can amplify selection on codons that become expressed in multiple contexts without changing the number of nucleotides under selection. However, measuring selection within OvRFs is a difficult problem because the effect of a nucleotide substitution (*i.e*., whether synonymous or non-synonymous) depends on multiple codon contexts (Pedersen and Jensen, 2001).

Computer simulations of biological processes are widely used as a tool to characterize biological processes that are otherwise too complex to represent as a mathematical model for analysis (Arenas, 2012). There is a great diversity of programs that simulate molecular evolution along a phylogeny (Rambaut and Grass, 1997), which are designed to model different aspects of evolution including recombination (Cartwright, 2005), insertions and deletions (Strope et al., 2007), variable selection intensities across the sequence (Hall, 2008), and different substitution biases (Spielman and Wilke, 2015). However, we have not found a publicly available program that can simulate evolution in linear or circularized genomes with an arbitrary distribution of overlapping and non-overlapping genes and variable codon-specific selection pressures, yielding a multiple sequence alignment for a given tree. A standard simplifying assumption in molecular evolution is that nucleotides or codon substitutions are independent and identically distributed outcomes of a continuous-time Markov model. However, a codon within an OvRF is no longer independently evolving, since a nucleotide substitution can change the selective context for subsequent changes at nucleotides of adjacent codons.

Here, we describe a simulation method (HexSE) implemented in Python that simulates molecular evolution along an input tree where the sequence may contain any number of OvRFs. We employ a memory-efficient data structure to track the rates of substitution events at every individual nucleotide of an evolving sequence.

## IMPLEMENTATION

### Input specification

HexSE takes three input files. First, it requires a FASTA- or Genbank-formatted file containing a nucleotide sequence (minimum 9 nt) to seed the simulation at the root of the tree. Next, the user must provide a configuration file in the form of a YAML-formatted file specifying the location of each ORF, including the parameters of the distribution that will be used to sample the frequency of mutations for each gene. Lastly, HexSE requires a file containing the Newick serialization of a phylogenetic tree. The tree must be rooted, *i.e*., contain a root node to assign the input sequence. Branch lengths of the tree are assumed to be scaled to the expected number of nucleotide substitutions per site, which is the standard unit for maximum likelihood reconstructions.

Additional parameters that may be specified in the YAML-formatted control file include: whether the genome is circular or not (default: false), a global transition/transversion ratio *κ* (default: 0.3); the parameters and number of rate categories for discretized gamma or lognormal distributions, which are used to model variation in selection (*ω*) and mutation rates (*μ*) for each reading frame; and the stationary nucleotide frequencies *π*. By default, *π* is set to the empirical frequencies in the root sequence.

### Algorithm

To simulate sequence evolution, we use a standard Gillespie (1976) algorithm for the exact stochastic simulation of discrete events. The model is initialized by assigning the input sequence to the root of the tree and calculating the rates of every possible substitution at position i to nucleotide *j*:

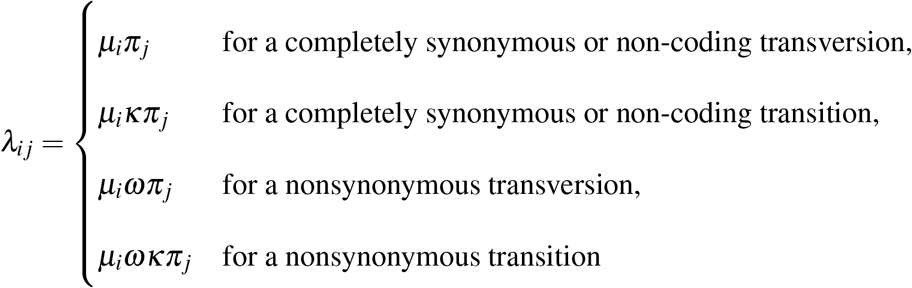

where *μ_i_* is the baseline mutation rate. The effect of selection is tracked by a vector **w** corresponding to the reading frames {−2, −1, −0, +0, +1, +2}, such that ||**w**|| = 6. An element of **w** is set to 1 if the substitution is synonymous in that reading frame, and the total selective effect on a substitution is 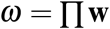. If a substitution is synonymous in all six reading frames, then *ω* = 1, *i.e*., the substitution evolves neutrally.

Next, we draw an exponentially-distributed waiting time to the next event, *t* ~ exp(-Λ), where 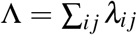. If *t* exceeds the length of the current branch, no event occurs and the current sequence is propagated to the next node. Otherwise, a substitution event is drawn with a probability proportional to the associated rate, *λ_ij_*/Λ, and the sequence is updated by that event. Ultimately, the substitution rates are re-calculated for the updated sequence. This process continues by level-order traversal of the phylogenetic tree until every terminal node (tip) of the tree carries an evolved sequence.

### Simulating a mutation

Computing and storing the rates of every possible substitution is time-consuming, and selecting the next substitution event by drawing a uniform random number would require a linear search of the cumulative probabilities. Since we use discretized probability distributions for *ω* and *μ*, these rates can only assume a finite number of values. Therefore, substitution events can be rapidly selected by traversing a hierarchical data structure that we call an event *probability tree*. All possible substitution events are stored at the tips of the event tree (Figure 2). Starting at the root of the event tree, we select the target nucleotide *j* with probability *π_j_*, moving down to the respective node at the next level. Next, we select the starting nucleotide *k* with probability 1/(1 + 2*κ*) if *k* → *j* is a transition and *κ*/(1 + 2*κ*) otherwise. We select the mutational rate category given the global rate distribution for *μ*, followed by selecting a particular region of the genome. When initializing the run parameters, we divide the sequence into categories determined by the distribution of open reading frames. For example, we divide the nucleotide sequence depicted in Figure 2 into five regions: nucleotides occurring within ORF *a* only (*p*1); ORF *b* only (p3); the overlapping region between ORF *b* and ORF *c* (*p*4); ORF *c* only (*p*5), and; nucleotides that are not in any ORF (*p*2).

**Figure 1:**
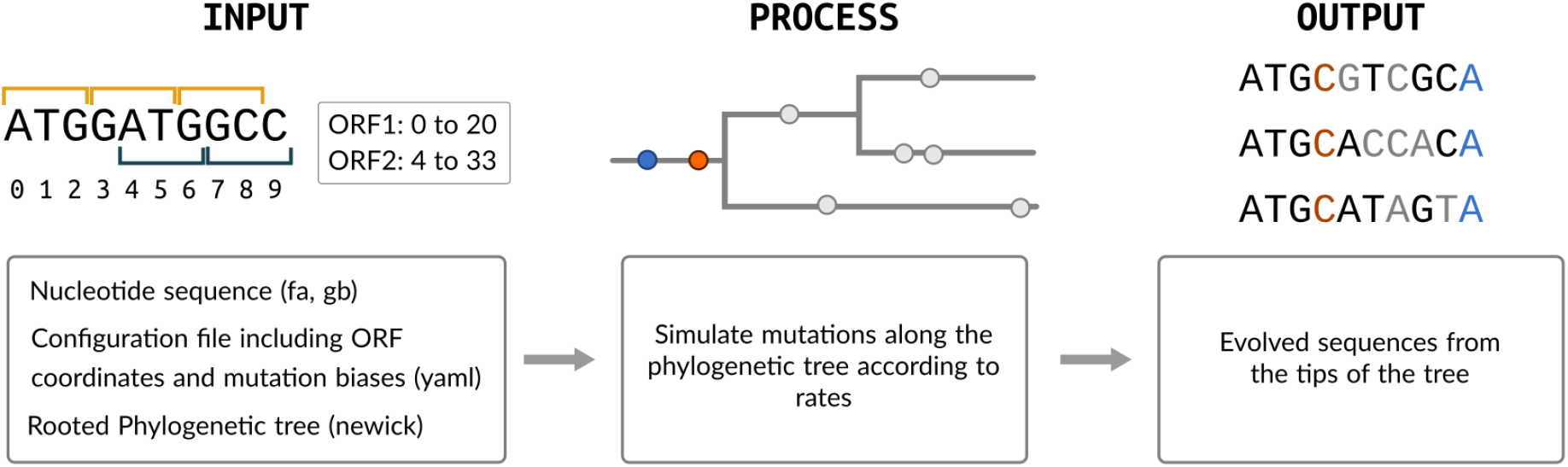
Pipeline overview. We use the Gillespie algorithm to simulate mutations along branches of the phylogenetic tree in order to create a nucleotide alignment.

**Figure 2:**
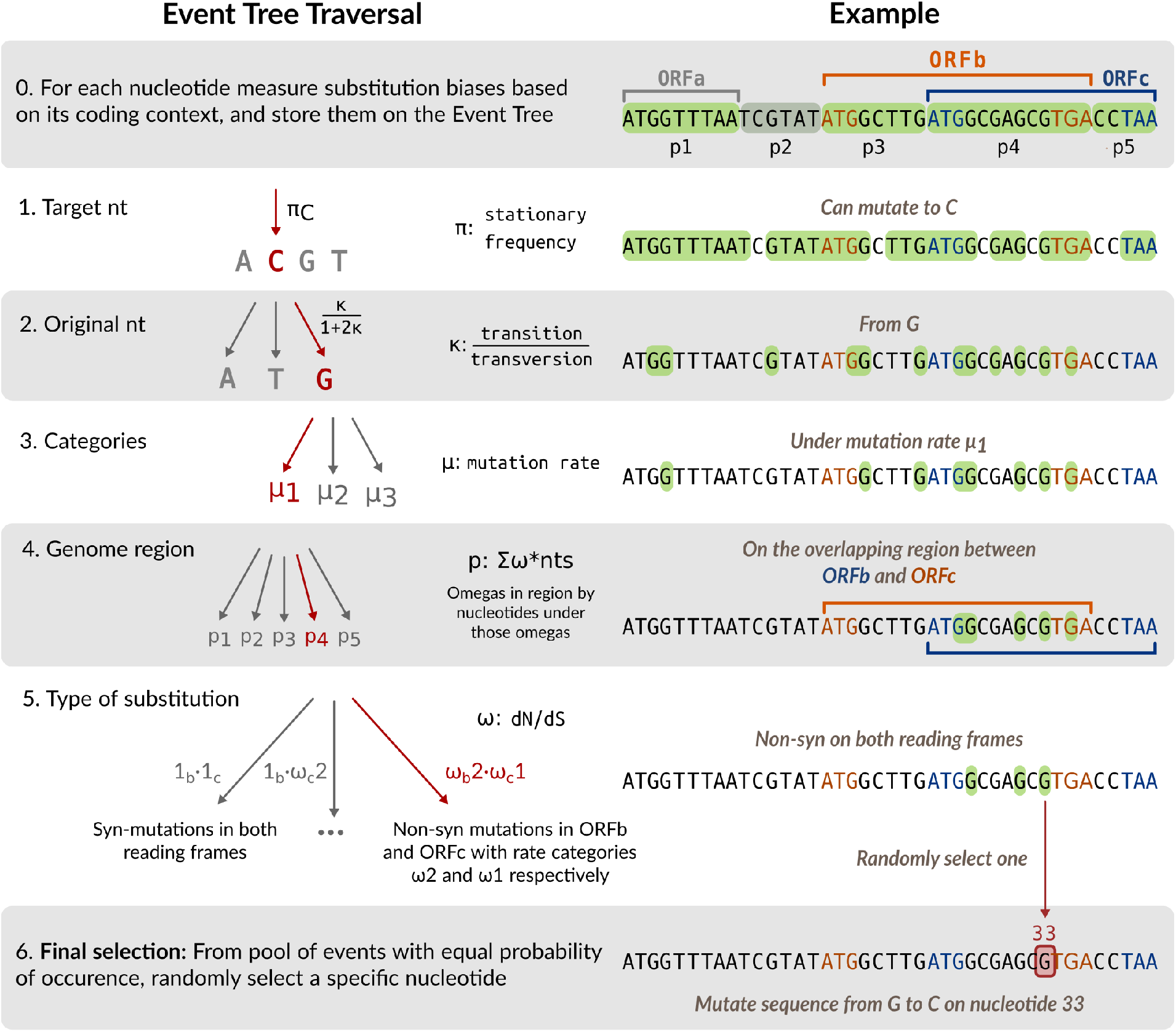
Traversal of an event probability tree to select mutations. An event probability tree is a data structure that we used to store nucleotide substitutions with the same probability. Each level of the event tree corresponds to an evolutionary parameter, such as the transition/transversion rate bias. Each branch represents a discrete value associated with the parameter represented by that level. Traversing the event tree from the root (left panel) selects progressively smaller subsets of mutations, as demonstrated in the right panel.

The probability of a region is proportional to the number of potential substitution events weighted by the net effect of selection, which is determined by the rate categories (distributions) of *ω*s associated with every ORF in the region. The weighted sum is calculated as 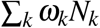, where *N_k_* is the number of nucleotide sites associated with the k-th combination of selection biases *ω_k_*, and the sum is computed over all such combinations in the region. For example, a substitution at position *i* that is non-synonymous in ORFs *a* and *b* with reading frames −1 and +0 can have a vector 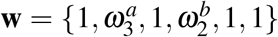, where 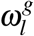 is drawn from a discretized distribution associated with ORF (gene) *g*. The value of *ω_k_* at position *i* results from multiplying the values assigned to 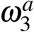 and 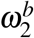.

Finally, we choose one of the *ω* combinations in the region, with a probability determined by the combination of selection rate biases. If a nucleotide is involved in a ‘start’ or ‘stop’ codon, then we do not include it in the event tree to prevent mutations from disrupting open reading frames.

Traversing the event probability tree from the root to a tip resolves the shared characteristics for a subset of substitution events. In other words, the tip stores references to the nucleotide substitution events that have the same probability of occurring. From this subset we can simply select an event uniformly at random, and update the evolving sequence accordingly. Since the composition of event subsets at the tips of the tree are updated with the sequence, we perform a deep copy of the event probability tree at every internal node of the phylogeny.

### Simulating evolution in the HBV genome

To demonstrate the usage of HexSE, we simulated evolution in the hepatitis B virus (HBV) genome, where around 30% of nucleotides within ORFs are involved in overlaps. Our inputs for the simulation were: the HBV reference genome (accession NC_003977.2) in Genbank format including coding sequence annotations for genes S, P, C and X; a random phylogenetic tree with 100 tips, using the implementation of the Kuhner and Felsenstein (1994) algorithm in T-REX (Boc et al., 2012); and the substitution model parameters. These parameters included a global mutation rate of 0.05, two rate categories for *μ* drawn from a lognormal distribution with two classes and shape 1.0 (resulting in rates 0.026 and 1.39 substitutions per nucleotide per unit time), a transition-transversion rate ratio of 0.3, and varying distribution parameters for *ω* in the four open reading frames. A second run was performed under the same parameters, but excluding all open reading frames except gene C (nucleotide coordinates 1,816 to 2,454).

Levels of genetic variation in simulation outputs varied substantially among codon sites, depending on their protein-coding context. We quantified these changes in genetic diversity using the Shannon entropy, 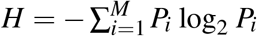, where *M* is the number of residues, and *P_i_* is the relative frequency of the *i*-th residue in the alignment column. This statistic provides a generic measure of diversity that applies to any combination of reading frames. We calculated the mean nucleotide entropy for sliding windows along the alignment with a width of 20 nucleotides and a step size of 1 nucleotide. For nucleotide residues, this statistic varies from zero, for a completely conserved site, to a maximum of *H* = 2.

A comparison of mean nucleotide entropy profiles between two simulation outputs on HBV sequences with one and four ORFs, respectively, is shown in Figure 3. In the simulation results where we included only one gene, we detected higher entropy in the non-coding regions (mean 0.539, IQR 0.475–0.598) which is consistent with the net effect of purifying selection in the coding region of gene C (mean 0.302, IQR 0.254–0.342). We observe similar mutation patterns in the non-overlapping region of gene C, on the alignment produced when all four ORFs are included. This region, comprised between nucleotides 1,840 and 2,309, has a mean of 0.306 (IQR 0.247–0.366), and reaches a minimum of 0.104. As expected, we noticed a significant differences in the entropy levels between overlapping and non-overlapping regions (average 0.304 vs 0.397; Wilcox rank sum test, *P* < 10^−15^). These shifts in mean entropy are also confounded by variation among ORFs in the intensity of purifying selection, as well as stochastic variation. These patterns are consistent with the cumulative effect of purifying selection in one or more ORFs on limiting diversification at the level of nucleotide sequences.

**Figure 3:**
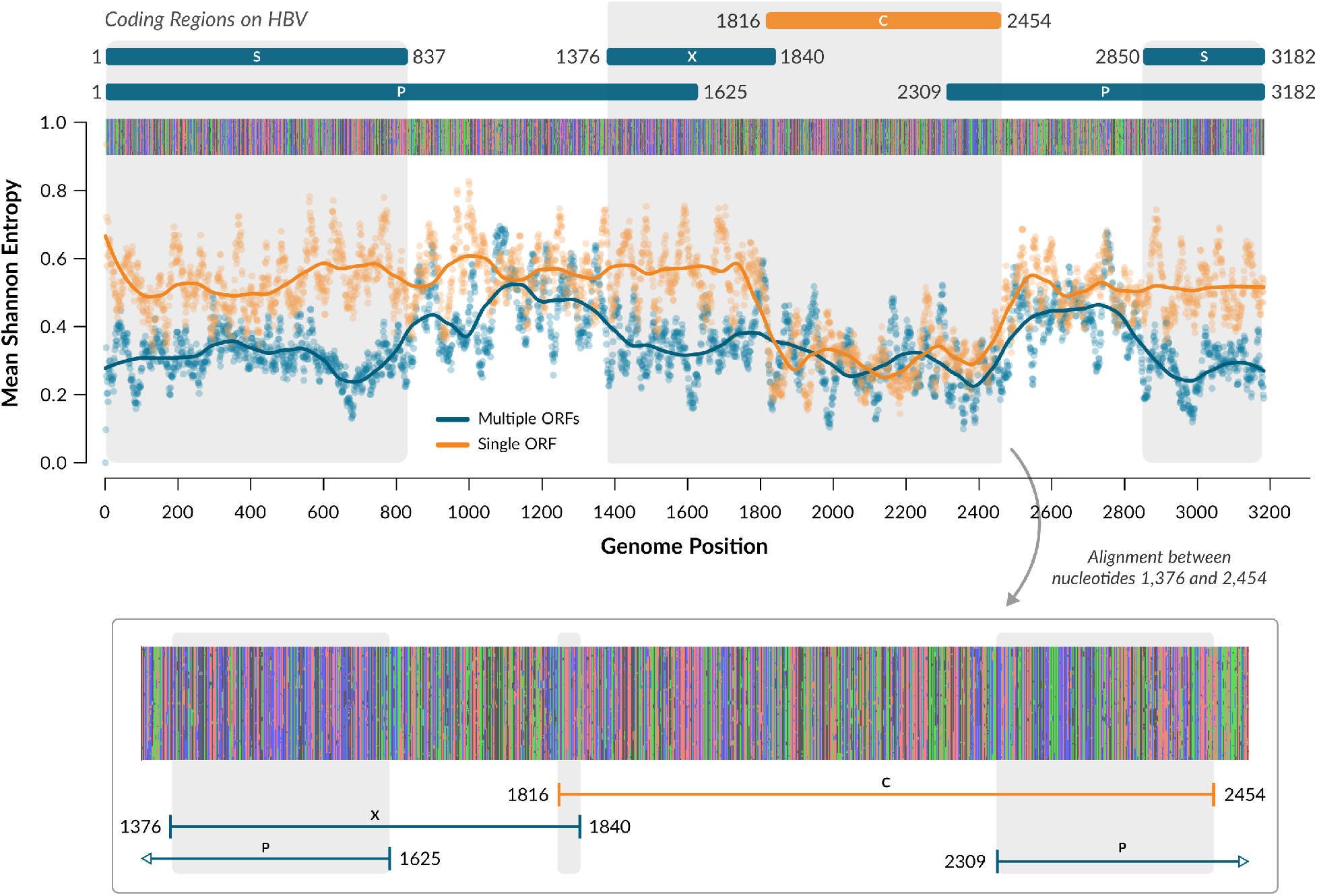
Simulating Evolution on HBV. Using HexSE, we simulated evolution in the HBV reference genome, with all four ORFs (blue) or only one ORF (orange). We calculated the nucleotide entropy per site for the resulting alignments to account for the number of mutations accumulated on each position over time. The region encoding gene C has similar entropy values for on both simulations. However, in the simulation results using the complete genome, the mean entropy falls from a mean of 0.306 to 0.222 in the overlapping region (nucleotides 2,309 to 2,454). There, mutations rarely accumulate, even in comparison with adjacent regions, which are also protein encoding sequences. In general, overlapping sites tend to accumulate less mutations than non-overlapping sites of the genome. This pattern is noticeable in the alignment between nucleotides 1,376 and 2,454, where three overlaps are present between genes X and P, C and X, and C and P.

The average time that HexSE requires to simulate evolution on HBV under our example parameters, is 52.74 seconds (interquartile range, IQR: 48.36–58.01s, *n =* 20) using four cores on an AMD Ryzen 5 3400g processor, with Python 3.6.9 and Ubuntu 18.04.6. In contrast, the same simulation takes about 310.3 seconds (IQR 281.3–308.1) to complete on a tree with 500 tips. In general, running times scale linearly with respect to the number of tips on the phylogenetic tree (Figure S1).

## ACKNOWLEDGEMENTS

This work was supported in part by a grant from the Natural Sciences and Engineering Research Council of Canada (NSERC Discovery Grant RGPIN 05516-2018).

## SUPPORTING INFORMATION

**S1 Figure.:**
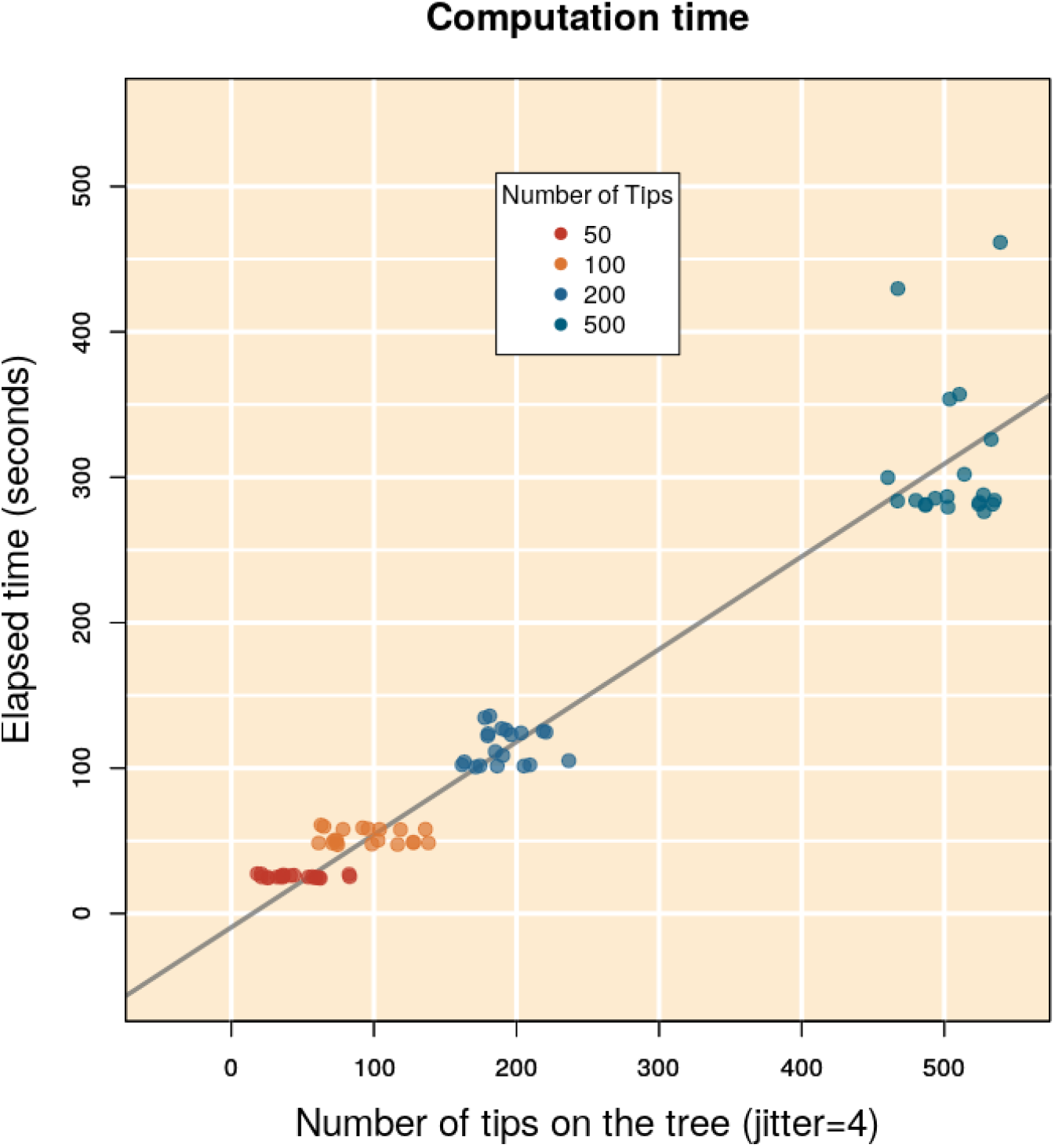
Elapsed times for 20 replicates of HexSE simulation runs on the HBV genome (accession NC_003977.2) using random phylogenetic trees with 50, 100, 200 and 500 tips. All simulations used the same parameters described in our example: global mutation rate of 0.05, two rate categories for *μ* drawn from a lognormal distribution with two classes and shape 1.0, and varying distribution parameters for *ω* in seven open reading frames.

